# Single acoustic closed loop stimulation in mice to modulate hippocampo-thalamo-cortical activity and performance

**DOI:** 10.1101/2022.04.16.488547

**Authors:** Sonat Aksamaz, Matthias Mölle, Akinola Esther Olubukola, Maxim Bazhenov, Lisa Marshall

## Abstract

Neural brain rhythms of sleep reflect neuronal activity underlying sleep-associated memory consolidation. The modulation of brain rhythms, for instance the sleep slow oscillation (SO) is used both to investigate neurophysiological mechanisms as well as to measure the impact on presumed functional correlates. In humans, auditory closed-loop stimulation targeted to the SO Up-state successfully enhanced the slow oscillation rhythm and phase-dependent spindle activity, although effects on memory retention have varied. Here, we aim to disclose relations between stimulation induced hippocampo-thalamo-cortical activity and retention performance on a hippocampus dependent task in mice. Closed-loop acoustic stimuli applied during four SO phases always acutely increased sharp wave ripple (SPWR) activity without disrupting non-rapid eye movement (NREM) sleep. Stimulation achieved an above chance preference index for stimuli delivered across a 3 h retention interval of sleep at the SO Up-state and at the Down-to-Up-state, but not at the Down-state nor late Up-state/Up-to Down-state. Results support the use of closed-loop acoustic stimulation in mice to investigate the inter-regional mechanisms underlying memory consolidation.

## Introduction

Sleep contributes or essentially facilitates memory consolidation (Stickgold 2005; Rasch and Born 2013; Tononi and Cirelli 2014). Reactivation and active systems memory consolidation, with a transfer of memory traces between the hippocampus and neocortex takes place (Rasch and Born 2013; Helfrich et al. 2019; Oyanedel et al. 2020), most notably for hippocampus-dependent, but also for long-term non-hippocampus dependent memories (Sawangjit et al. 2018; Schapiro et al. 2019). The temporal coordination of cortical slow oscillations (SOs), thalamo-cortical sleep spindles and hippocampal sharp-wave ripples (SPWR) reflects the neuronal communication between the underlying structures.

A method to investigate memory functions associated with brain rhythms during sleep is through their interrogation (Campos-Beltran and Marshall 2017; Malkani and Zee 2020). For example in rodents, enhanced coupling between sharp-wave ripples and delta spindles achieved by timed electrical stimulation enabled successful consolidation of weak memories (Maingret et al. 2016). On the other hand, reduced spindle-ripple coupling and impaired memory resulted from targeted disruption of spindle activity (Novitskaya et al. 2016; Swift et al. 2018). Similarly, suppression of hippocampal ripples was detrimental on memory (Girardeau et al. 2009; Ego-Stengel and Wilson 2010), indicating the relevance of each of these rhythms and their temporal interplay for memory consolidation.

In humans multiple studies have employed noninvasive weak electric and sensory stimulation procedures to investigate relations between brain rhythms and memory function, and to probe for future clinical applications (Marshall et al. 2006; Ngo et al. 2013b; Ketz et al. 2018; Salfi et al. 2020; Ladenbauer et al. 2021). Acoustic closed-loop stimulation (ACLS), in which acoustic stimuli was applied during two successive SO Up-states has proven robust in modulating endogenous SO and spindle activity in humans (Ngo et al. 2013b; Ong et al. 2016), although cognitive benefits are not straightforward (Henin et al. 2019; Schneider et al. 2020; Harrington et al. 2021). Most recently, a single closed loop acoustic stimulation during NREM sleep in rats affected EEG spectral slow wave and spindle power, and impaired learning performance on a non-hippocampus dependent task, when delivered at the SO Down-state (Moreira et al. 2021). Recent modeling study revealed specific impact of the acoustic stimulation when stimuli were delivered near Down-to-Up transition of SO (Wei et al. 2020).

Acoustic stimuli directly evoke both thalamo-cortical and hippocampal responses yet the interactions between responses to SO ACLS and memory formation are undisclosed. The aim of the current study was therefore to investigate the thalamo-cortical and hippocampal interactions induced by SO ACLS, and the behavioral impact. Since responses of the auditory system reveal stimulus and species specific-differences, even between rodents (Süer et al. 2004; Cromwell et al. 2008; Ngo et al. 2013a; Ngo et al. 2015; Witten et al. 2016), the present study investigated electrophysiological and behavioral response to a single acoustic stimulus given in a closed loop fashion dependent upon a detected SO of NREM sleep (sACLS). Retention performance was assessed on the hippocampus-dependent object place recognition (OPR) task (Mumby et al. 2002; Barker and Warburton 2011). We expected increased retention for stimulation around SO Up-state and for this to coincide with indications of increased inter-regional neural communication during sleep. We found increased retention performance for sACLS at the SO Up-state to be weakly related to assessed inter-regional neural communication at the beginning of NREM sleep, but an indication toward a role of preREM sleep. Moreover, acoustic stimuli at all SO phases produced a post-stimulation suppression of spindle activity in the second range.

## Results

### A preference index above chance is not maintained for sACLS at 2 and 300 ms after the SO positive half-wave

To investigate the modulatory effects of sACLS on sleep related brain rhythms and on memory retention, mice were subjected to five sessions of an OPR task with sACLS at the beginning of their inactive phase. During the 3 h interval between OPR sample and test phases, in separate sessions sACLS was applied at four different delay times (2ms, 120ms, 180ms, 300ms after the positive SO half-wave peak, i.e., Down-state) during NREM sleep and with sham stimulation (cp. **Figure 1)**. The delays correspond to the SO Down-state, Down-to-Up-state transition, Up-state, and late Up-state/Up-to-Down-state transition. The sham session was devoid of acoustic stimulation, yet triggers were set at online SO detected SOs as in the other sessions (**Figure 1A, 1B**).

**Figure 1.**
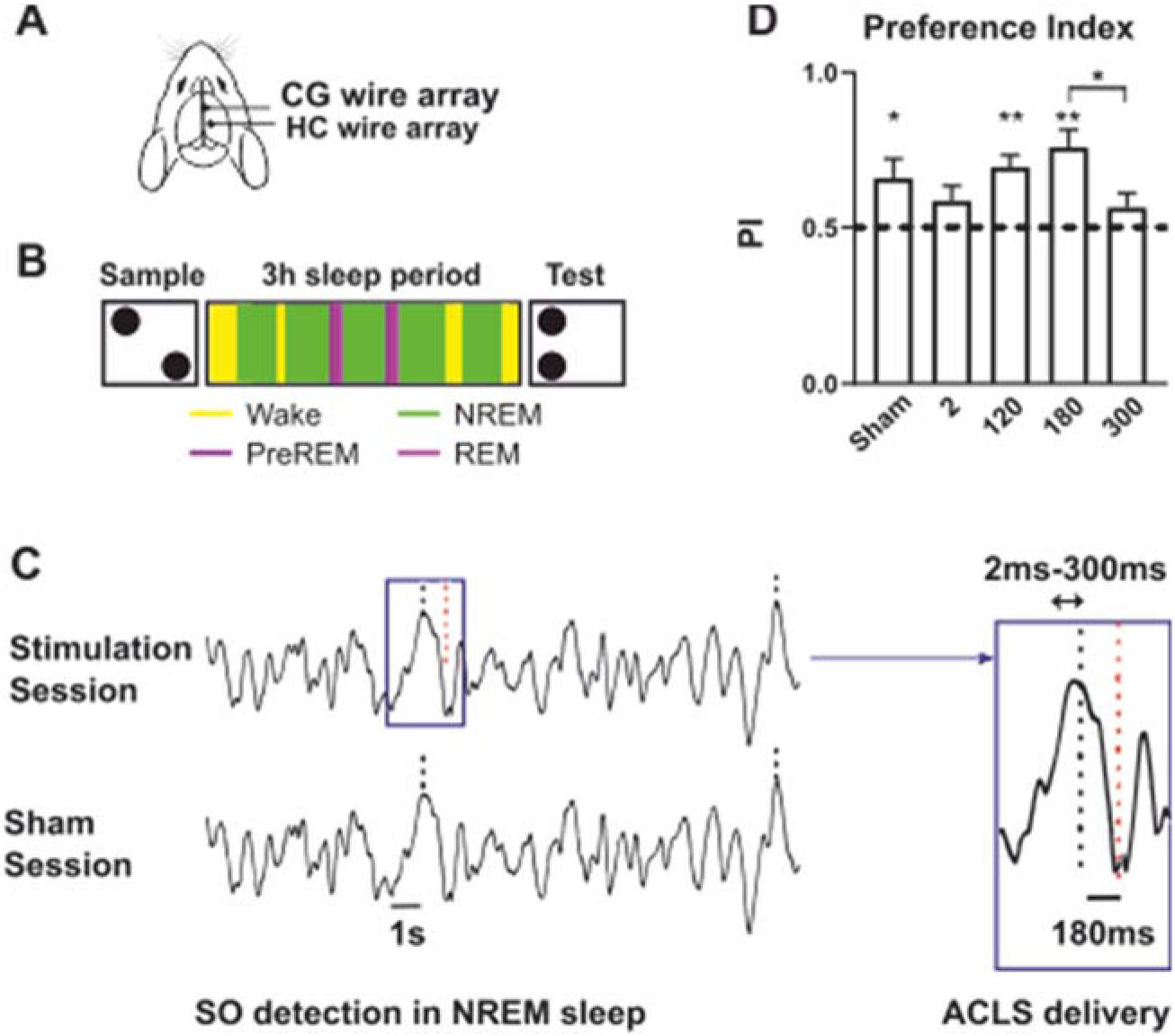
Visualization of electrode positons, experimental design, sACLS application, and behavioral results, e. **A.** Positions of cortical and dorsal hippocampal wire array electrodes on the mouse brain**. B.** Time frame of a daily OPR session. Following the lights on, the animals were placed in the open field for 10 min to perform the sample phase of the OPR test. Thereafter animals were tethered to the amplifier and allowed to sleep for 3h in the recording. After the sleep period, animals were untethered and transferred to the open field for the OPR test phase (5min). **C.** SO detection and sACLS application. Left: Stimulation session, online SO detection (black dashed line) followed by acoustic stimulation (red dashed line). Subsequent pseudo-stimulation after a minimal inter-stimulus interval of 2.5 s is not shown. In the sham session SO detection remained without an acoustic stimulation. Right: SO up-state sACLS at 180 ms delay is shown as an example. **D.** Mean (± SEM) preference for the replaced object across all stimulation conditions. Deviation from the chance level (dotted line) was represented with stars, significant difference between conditions were shown with asterisks. Number of animals in each condition: sham, 2ms, 180 ms, n=10; 120 ms, n=7; 300 ms, n= 6. * p < 0.05, **p < 0.01.

Our choice of a 3 h delay period was based on previous reports on object place recognition tasks in mice with a shorter delay period (Dere et al. 2005). We thus hypothesized that a preference index significantly above chance would not be obtained with a 3 h delay period in sham but would be obtained by sACLS given at an optimal delay. Mice performed above chance levels for sACLS at 180ms (Up-state), at 120ms (Down-to-Up-state-transition) as well as in the sham session (**Figure 1D**, one sample t-test; theoretical mean: 0.5, 180 ms: t(9) = 4.613, p=0.001; 120 ms: t(6) = 5.039, p=0.024; sham: t(9) = 2.525 p = 0.033). However, for sACLS delivered at 2 ms (Down-state) or at 300ms (Up-to-Down-transition), the preference index failed to surpass the chance level: 2 ms, t(9) = 1.784, p=0.1081; 300 ms, t(5) = 1.375, p=0.2275). Although across all conditions a trend was found (mixed-effects model; F (4, 38) = 2.105, p = 0.099) we proceeded to test according to our hypothesis. Paired comparisons between conditions support increased retention performance for sACLS at the 180ms Up-state, but also for the preceding Down-to-Up transition at 120 ms (180ms vs. 300ms, (t (5) = 3.362, p = 0.020, paired t-Test); 180 vs. 2 ms t (9) = 2.164, p= 0.059; 120 vs. 300 ms, t (5) = 2.113, p = 0.088).

### Event related ripple RMS at 180 ms overlaps with the endogenous increase in ripple RMS

First, we calculated the event related responses of the main brain rhythms to sACLS. **Figure 2.** reveals the mean changes in local field potentials (LFP) and root mean square (RMS) ±1 s around the corresponding acoustic or pseudo-stimulation for the 3h-interval. Event related LFPs at the cingulate cortex confirm that stimulation adequately targeted the four different SO phases (**Figure 2**). Event related spindle RMS increased dramatically on stimulation, however, after peaking around the time point of acoustic stimulation for all sACLS delays (**Figure 2A)**, instead of maintaining a heightened level spindle RMS decreased around 120 ms after sACLS and maintained a low level (**Figure 2B**).

**Figure 2.**
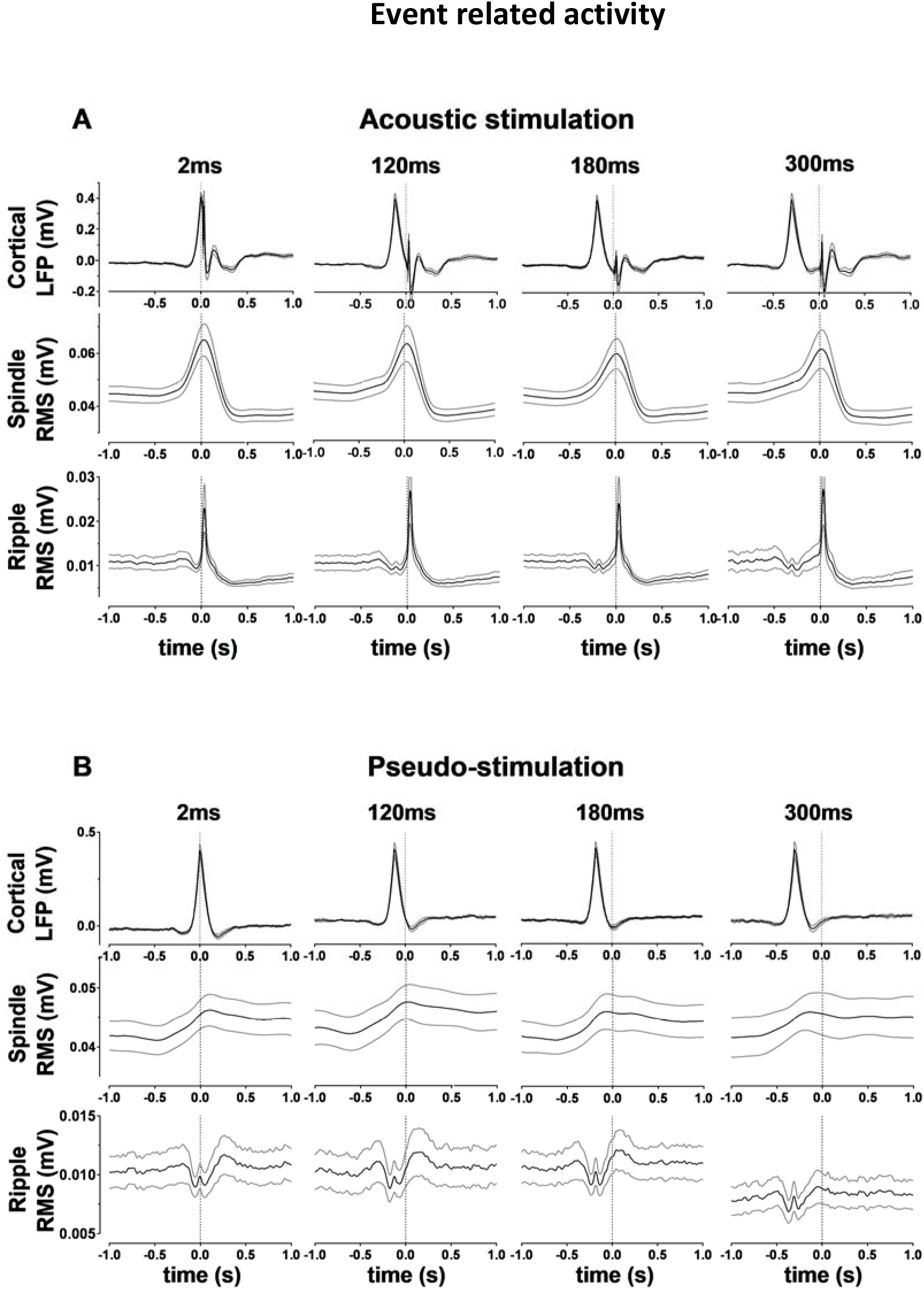
sACLS delivery time-locked to the differential SO-phases modulates cortical local field potential (LFP), cortical spindle and hippocampal ripple activity. **A.** Acoustic stimulation. **B.** Pseudo-stimulation. **A, B.** From top to bottom, mean (± SEM) cortical LFP, average spindle RMS, and average RMS of CA1 SPWRs within ±1s of stimulation. 2ms, 180 ms, n=10; 120 ms, n=7; 300 ms, n= 6.

Endogenous ripple RMS reveals the unique temporal pattern (**Figure 2B**) of decreasing at the Up-to-Down state, revealing a small, short-lived peak around the Down-state, and increasing gradually with the SO Down-to-Up transition, reaching maximal mean values around the time of the endogenous SO Up-state. **Figure 2A** reveals that this pattern is completely disrupted at 2ms sACLS, at 120 ms delay it is recognizable, and at 180 ms the evoked ripple RMS overlaps in time with the endogenous ripple RMS maximum, which is visible for pseudo-stimulation. At 300 ms delay evoked ripple RMS occurs after the time of the endogenous maximum. The increased variability at 300 ms sACLS is unlikely only due to the lower animal count as subject numbers were similar at 120 ms sACLS.

### sACLS application did not disrupt NREM sleep

To control whether sACLS may have modified sleep composition, we investigated the proportion of sleep stages within the 3h post-learning interval across differential stimulation conditions (**Table 1A**). Only the duration of PreREM sleep revealed an effect of Condition (F (4, 29) = 2.911, p = 0.039). PreREM sleep duration at the 180 ms delay was longer than that at 300 ms (mean difference 2.46 ± 0.80 min, p=0.044, post-hoc Holm-Sidak multiple comparison test). No other sleep stage differed in duration between conditions (Table 1.A).

**Table 1.**
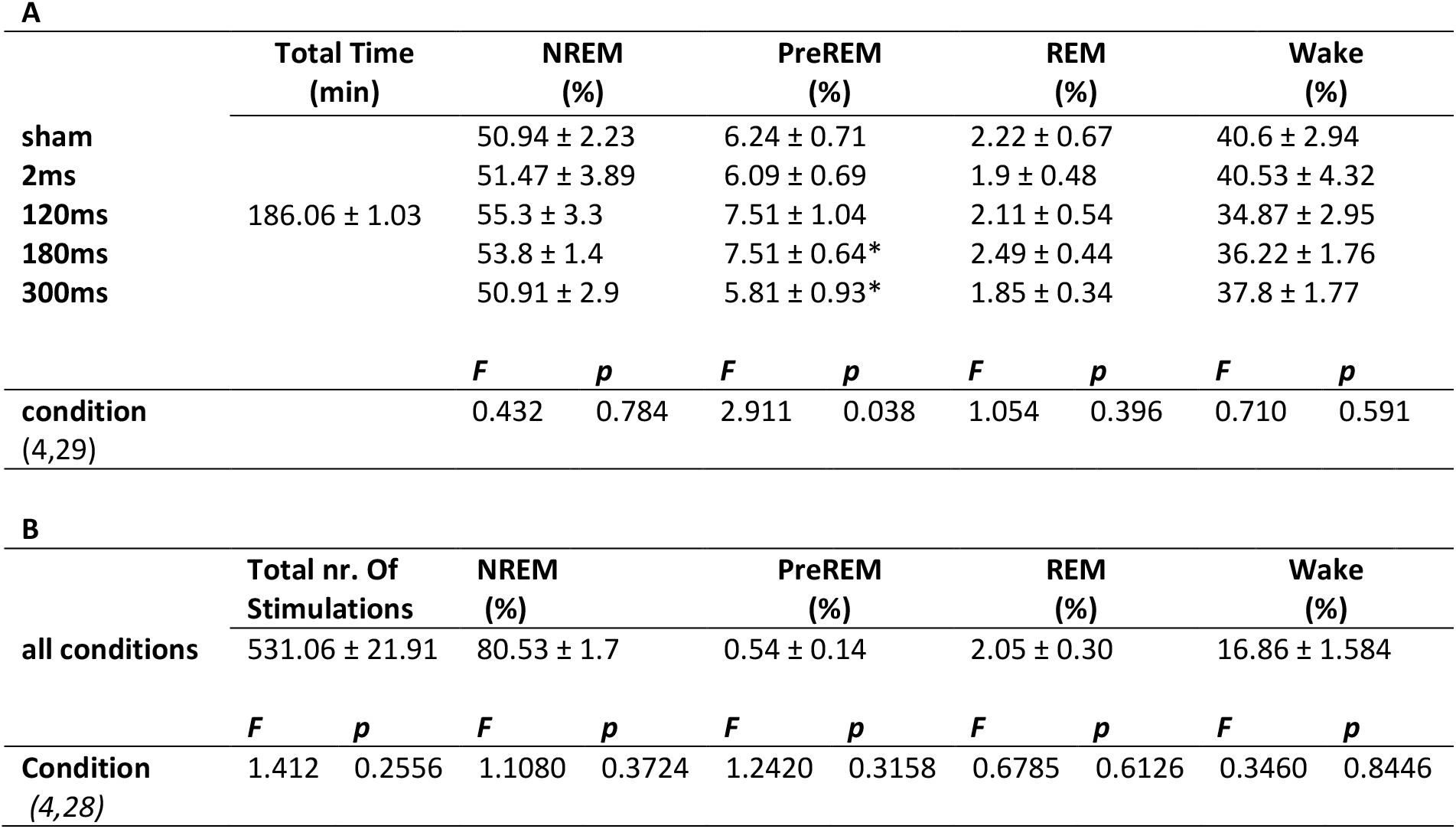
sACLS did not disrupt NREM sleep. **A.** Distribution of sleep stages during the 3h recording interval. * p < 0.05, for the difference between the 180ms and 300ms sessions for PreREM sleep duration, post-hoc Holm-Sidak multiple comparison test. **B.** Distribution of acoustic stimulations during the respective sleep stages as percent of the 3h recording interval, averaged across conditions. F and p values of mixed effects models are given. Number of animals in each group: 2ms, 180 ms, n = 10; 120 ms, n = 7; 300 ms; n = 6; sham, n = 9. In the sham session triggers of one animal could not be retrieved. Mean ± SEM

As shown in **Table 1B,** neither the total number of acoustic stimulations per condition nor the distribution within any of the sleep stages differed. The unexpected large number of stimulations during wakefulness for all conditions is most likely due to a poor online distinction between NREM sleep and quiet wakefulness in some animals in which poor EMG recording quality did not permit optimal assessment of muscle tone.

### sACLS did not affect individual oscillatory sleep events

Following the affirmation that NREM sleep duration was preserved across the different sACLS conditions, sleep-event parameters relevant for memory retention were analyzed. Neither density, amplitude nor duration of SOs, spindles nor SPWRs revealed a significant Condition effect (each p ≥ 0.1, mixed-effects analyses), although spindle duration revealed a trend (F (4,27) = 2.136, p= 0.1037). Since duration of PreREM sleep, during which spindle-like activity occurs, differed between 300 and 180 ms, we specifically compared corresponding spindle durations. Mean spindle duration was longer in the 180 ms than 300 ms delay condition (t(6)= 2.585, p= 0.049, uncorrected for multiple comparisons).

Furthermore, neither SO slope, nor spindle and ripple RMS revealed a significant effect of Condition (each p ≥ 0.1, mixed-effects analyses). Values presented in **Table 2** are thus mean values collapsed across all conditions. Results on sleep-event parameters per condition are reported below for early and late NREM sleep.

**Table 2.** Average sleep-event parameters across sACLS conditions. Event parameters of density, amplitude, duration, and slope or RMS, respectively were analyzed for SO, spindle and SPWR events within NREM sleep of the 3h post-learning interval. Mean ± SEM are given for all groups independent of the condition. SO and ripple parameters, n = 43; spindle parameters, n = 41. Spindle density values of one animal were spuriously high, thus data of this animal were omitted.

| Event | Density ($\text{min}^{-1}$) | Amplitude (mV) | Duration (ms) | RMS ($\mu\text{V}$) | Slope (mV/s) |
| --- | --- | --- | --- | --- | --- |
| SO | $12.32 \pm 0.69$ | $0.62 \pm 0.03$ | $703 \pm 4.14$ | - | $0.83 \pm 0.04$ |
| Spindle | $2.36 \pm 0.15$ | $0.40 \pm 0.011$ | $843 \pm 13.5$ | $93 \pm 2.55$ | - |
| SWPR | $15.60 \pm 1.13$ | $0.18 \pm 0.010$ | $70 \pm 0.75$ | $39 \pm 2.19$ | - |

### Sustained suppression of spindle and ripple event-activity following sACLS

Spindle and ripple RMS already indicated specific sACLS induced temporal response dynamics (**Figure 2**). Thus, we next compared SO, spindle and SPWR event-activity in response to sACLS directly between acoustic and pseudo-stimulation (**Figure 3**, Supplemental_Fig_S1.pdf). Expectedly, SO event-activity was larger for both acoustic and pseudo-stimulation around 0s corresponding to the detected SO positive half wave peaks. Notably, acoustic stimulation at the Down-state (2ms delay condition) strongly suppressed SO activity within −0.1s to 0.1s peristimulus. Suppression of SO event-activity at the 120 and 300 ms delays was shorter and lower (**Supplemental_Fig_S1.pdf)**. Significant suppression did not occur at the 180 ms delay. SO event-activity was increased following acoustic stimulation in the 2 ms delay (0.5-1.0 sec range), whereas in the 300 ms delay condition SO activity was subsequently suppressed (0.5-0.6 s range; **Figure 3A, Supplemental_Fig_S1.pdf**).

**Figure 3.**
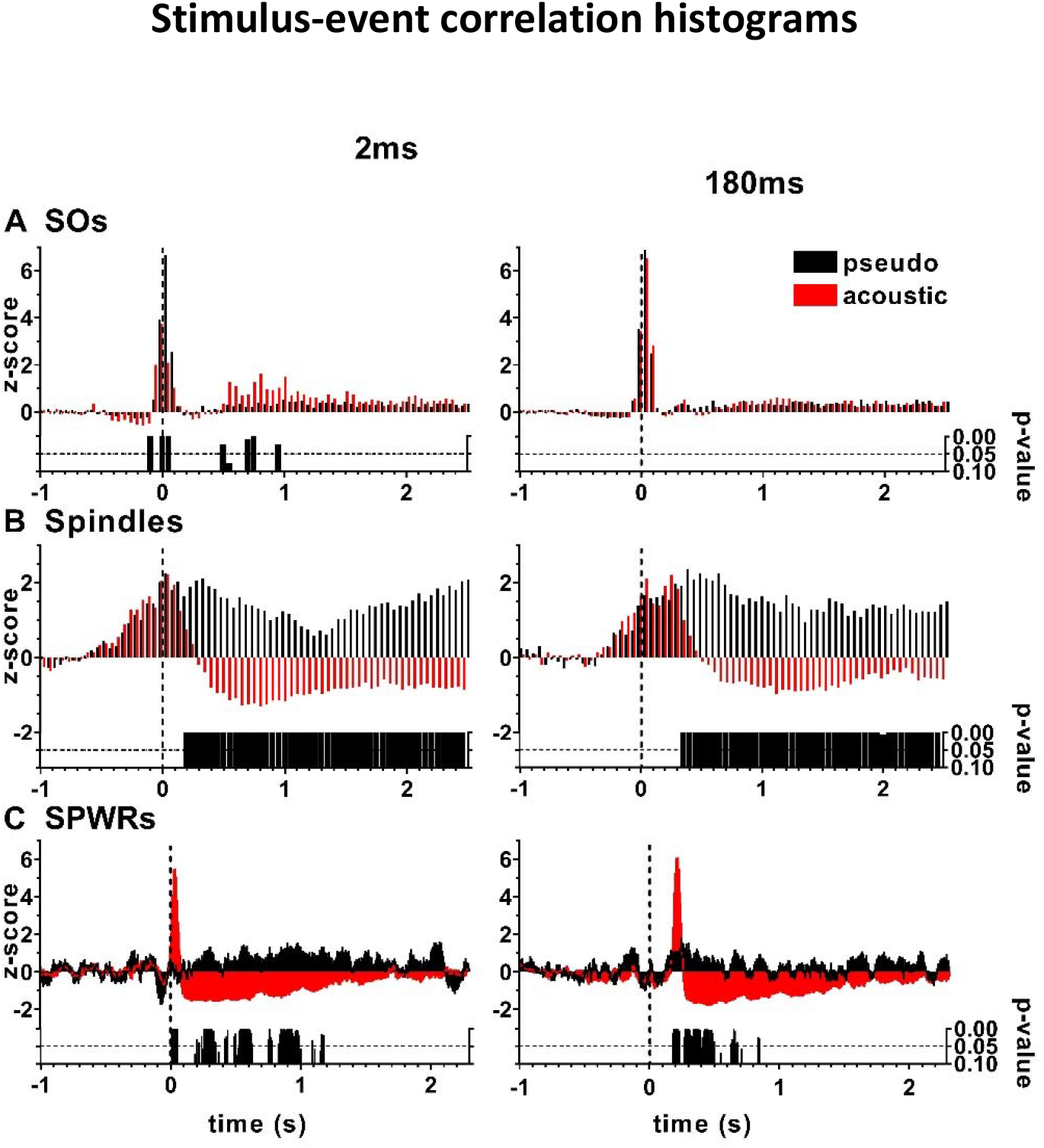
SO, thalamo-cortical spindle and hippocampal ripple event-activity relative to sACLS. **A-C.** Mean stimulus-event correlations for SO, spindle and CA1 SPWR events (z-transformed) for acoustic and pseudo-stimulation averaged across the 3h interval for the 2 ms and 180 ms delay conditions, respectively. T = 0 corresponds to the detected SO Down state peak. Data are baseline normalized across the first 0.5 seconds. Bar charts depict the Sidak-Holms adjusted bin-wise p-values of two-tailed t-tests. Results are presented for the two delay conditions with behavioral preference index above (180ms) and below chance level (2ms), and for which data of all ten animals are available. Comparable figures for the other two delays are given in the **Supplemental_Fig_S1.pdf.**

For spindles stimulus-event correlations show highest event-activity around the time of acoustic stimulation. Subsequently, with varying latencies spindle stimulus-event activity was consistently suppressed for at least 1.5 s by the acoustic as compared to pseudo-stimulation (**Figure 3B, Supplemental_Fig_S1.pdf**). As already noticeable in Figure 2, hippocampal SPWR stimulus-event correlations increased drastically with acoustic stimulation as compared to pseudo-stimulation. Stimulus-event correlation histograms also revealed a suppression of SPWR event-activity after acoustic stimulation, but only up to ~ 1 s (**Figure 3C., Supplemental_Fig_S1.pdf**).

Together, acoustic stimulation at all delays led to a sharp increment in ripple stimulus-event correlations, and post-stimulation attenuation of both spindle and SPWR event-activity. Alone for the 180 ms delay acoustic stimulation remained without an effect on SO stimulus-event correlations as compared to pseudo-stimulation.

### Is retention performance reflected by enhanced coupling to SO activity?

A prominent role of SOs in promoting memory consolidation is to group other sleep events. Therefore, we investigated if sACLS application at the different SO phases affected the coupling between SO, thalamo-cortical spindle and hippocampal ripple activities. Here offline detected SOs of NREM sleep throughout the entire 3 h sleep interval were analyzed, i.e., independently of the stimulus-triggered responses as given in **Figures 2, 3**.

All event-event correlation histograms of sham reflect the characteristic temporal relationships between the neural oscillations: Spindle – SO event-event correlations are minimal at the SO Down-state and increase strongly at the Down-to-Up transition (Sham in **Figure 4A**). Similarly, ripple event-event correlations are minimal around the SO Down-state, increase at the Down-to-Up transition and peak around the Up-state (sham in **Figure 4B**). Regarding temporal coupling of ripple to spindle event-activity (sham in **Figure 4C)**, the former is maximal around the deepest spindle trough, i.e., around spindle-midpoint. Vice versa, coupling of spindle to ripple event-activity (**Figure 4D**) shows a broad surge in spindle activity around the deepest trough of hippocampal SPWRs.

**Figure 4.**
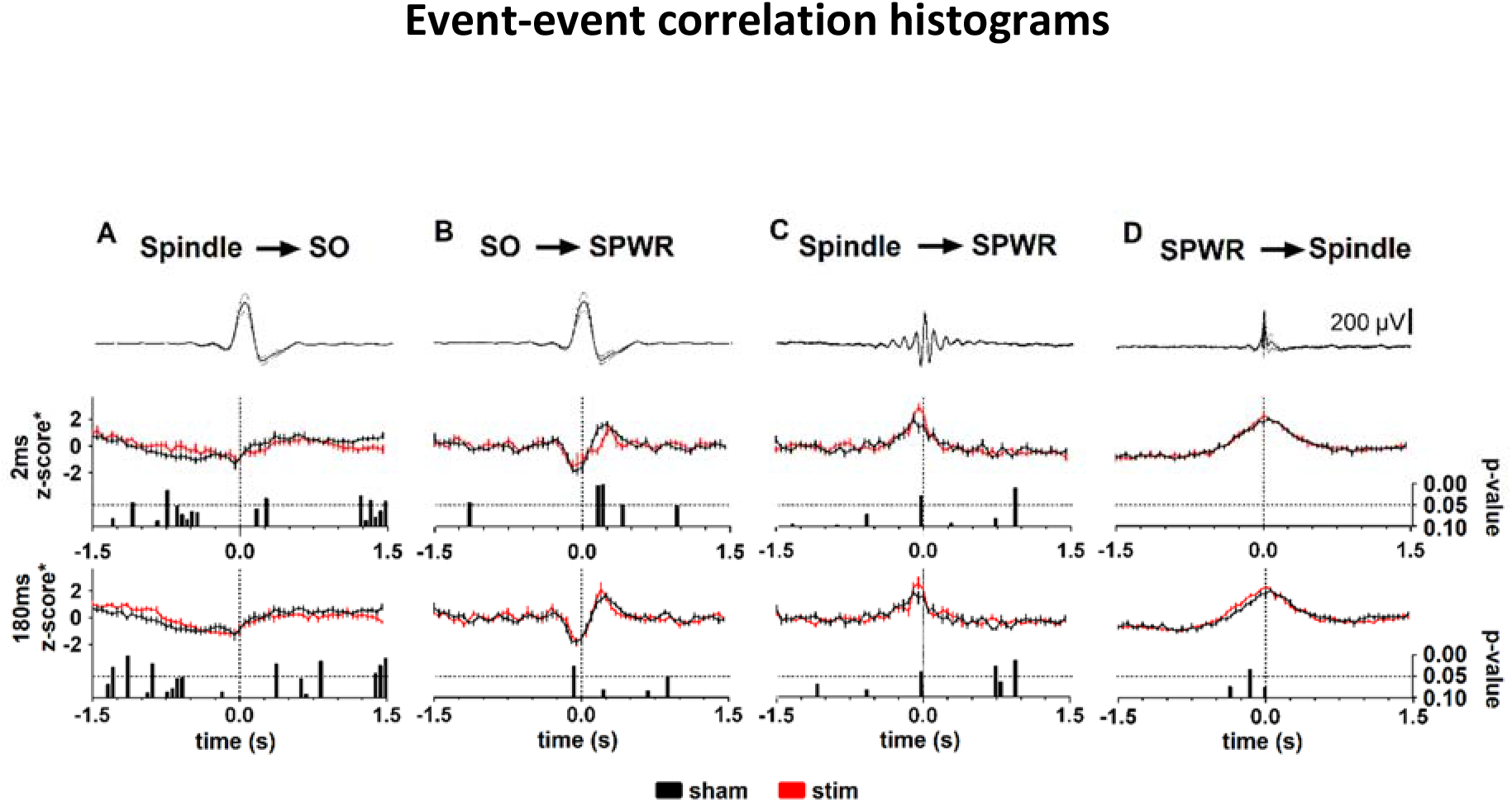
Event-coupling histograms between SO, spindle and SPWR event activity, respectively, across the 3h interval. **A.** Spindle event-activity time-locked to the positive half-wave peak of the SO (t= 0). **B.** SPWR event-activity time-locked to the positive half-wave peak of the SO (t= 0). **C.** Ripple event-activity time-locked to the deepest trough of the cortical spindle. **D.** Spindle event-activity time-locked to the deepest trough of hippocampal ripples. **A-D.** Upper diagrams represent the mean reference event in sham condition, bottom bar charts depict the bin-wise p values of two-tailed t-tests (not corrected for multiple comparisons). Mean ± SEM. Results are presented for the two delay conditions with behavioral preference index above (180ms) and below chance level (2ms), and for which data of all ten animals are available. Comparable figures for the other two delays are given in the **Supplemental_Fig_S2.pdf.**

Unexpectedly, for all stimulation conditions event-event correlation histograms are largely similar to sham. At most, a decrease in temporal coupling of ripple event-activity to SO activity around 0.25 s stands out for the 2 ms delay (**Figure 4B**). This is similar for the 300 ms delay regarding this time window, but not observed at 180 ms sACLS (**Supplemental_Fig_S2B.pdf**).

### Focus on early NREM sleep

Since increased hippocampal reactivation and increased spindle density within the first 1-2 hours of sleep are reported (Gais et al. 2002; Eschenko et al. 2006; Sawangjit et al. 2018; Giri et al. 2019), we conducted analyses separately for early and late sleep periods. Early and late sleep periods contained closely similar amounts of NREM sleep (early/late NREM: sleep 48.3 ± 1.3 min and 47.9 ± 1.5 min, respectively; F (1, 9) = 9.5e-005, p = 0.992, for the main effect of Time, mixed-effects model). Number of stimulations per sleep period also did not differ (F (4, 36) = 0.915, p = 0.466 for the main effect of Condition; (F (4, 22) = 1.279, p = 0.308 for the interaction Condition x Time).

Distribution of REM and preREM sleep were as anticipated. Durations of both were longer in the late sleep period (REM sleep: F (1, 9) = 21.76, p= 0.0012; preREM sleep F (1, 9) = 67.01, p <0.0001, for the main effect of Time). As reported for the 3 h interval, preREM sleep revealed a significant Condition effect (F(4,36) = 4.009, p = 0.0086). There was no significant Condition x Time interaction for REM nor for preREM sleep (preREM: F(4,22) = 1.352, p= 0.2825, REM: F(4,22) = 0.5954), nor a Condition effect for REM sleep (F(4,36) = 0.5952, p = 0.6684).

Sleep oscillatory events in NREM sleep also revealed characteristic temporal changes from early to late sleep periods consistent for all conditions. In early NREM sleep, SO amplitude, density and slope were higher and steeper, respectively (**Figure 5**). Across all conditions SO and spindle durations were shorter in early NREM sleep. However, as for the 3h interval, spindle duration tended to differ between conditions (F(4,36)=2.597, p =0.053). SPWR duration, density, amplitude and RMS were all higher in early NREM sleep. Values and statistics of oscillatory event parameters are given in **Supplementary Tables.2-4.**

**Figure 5.**
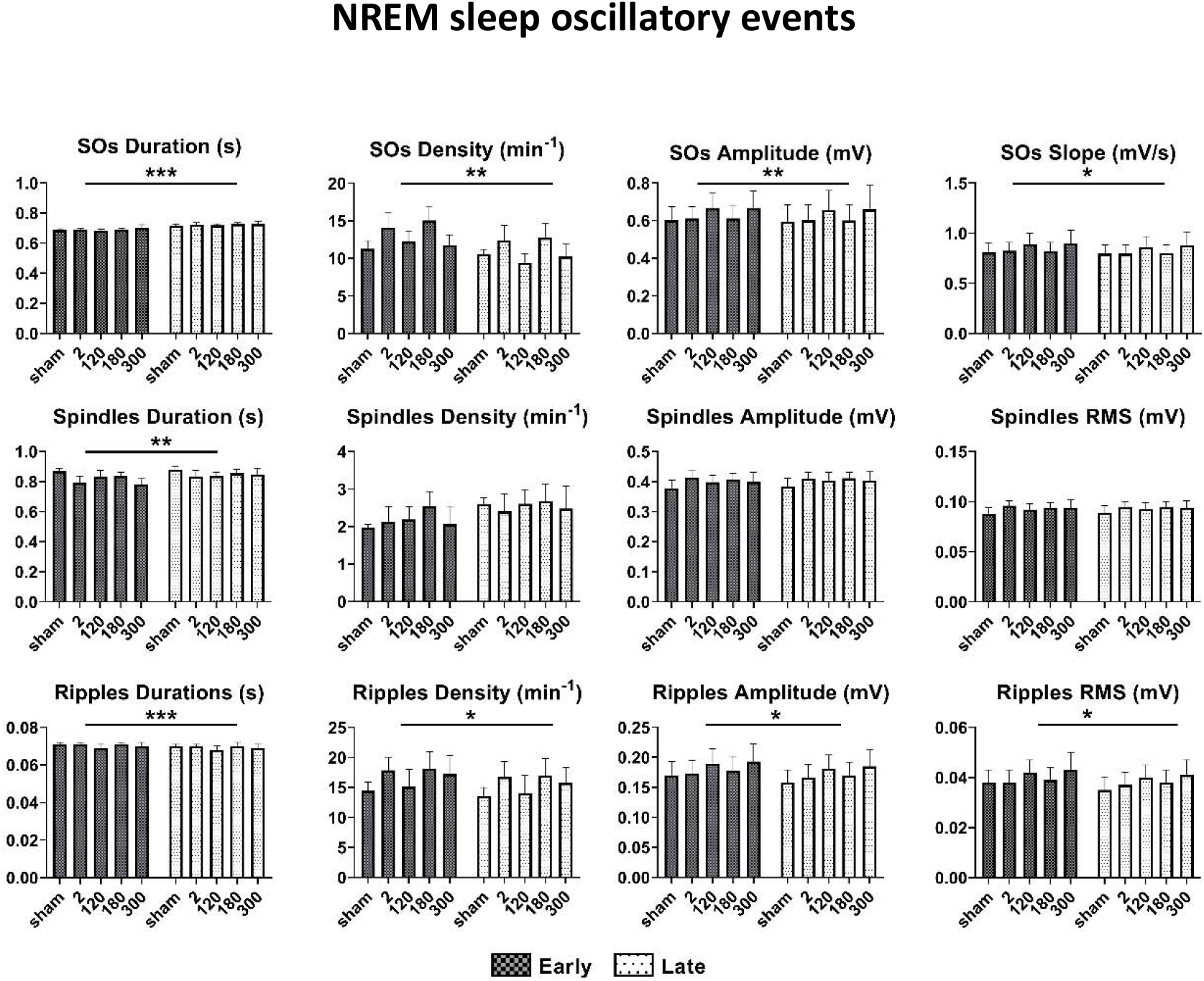
Sleep oscillatory event parameters for early and late NREM sleep. Mean (± SEM) SO, spindle and CA1 SPWR event parameters measured during early and late NREM sleep are compared. Durations of SO, spindle and SPWR oscillatory events correspond to the time from beginning to end of the detected oscillatory event. Density values apply to NREM sleep. Amplitude values represent peak-to-peak values of the corresponding filtered signal. Spindle and ripple RMS are derived from detected oscillatory events. SO slope is derived from the positive Down state peak to the following zero crossing. Asterisks indicate significant effects of Time. Error bars: * p < 0.05, **p < 0.01.

### SO-spindle coupling dynamics in early NREM sleep

Since a modification of SO-spindle coupling with acoustic closed loop stimulation is most pronounced, we were interested in whether effects of sACLS on coupling in early NREM sleep would be more pronounced. **Figure 6** suggests that at the 180 ms delay spindle event-activity was more strongly amplitude modulated at the SO Up-to-Down transition relative to sham (range, −0.3 to −0.1 s), whereas at the 2 ms condition modulation is decreased compared to sham (−0.75 to −0.35 s).

**Figure 6.**
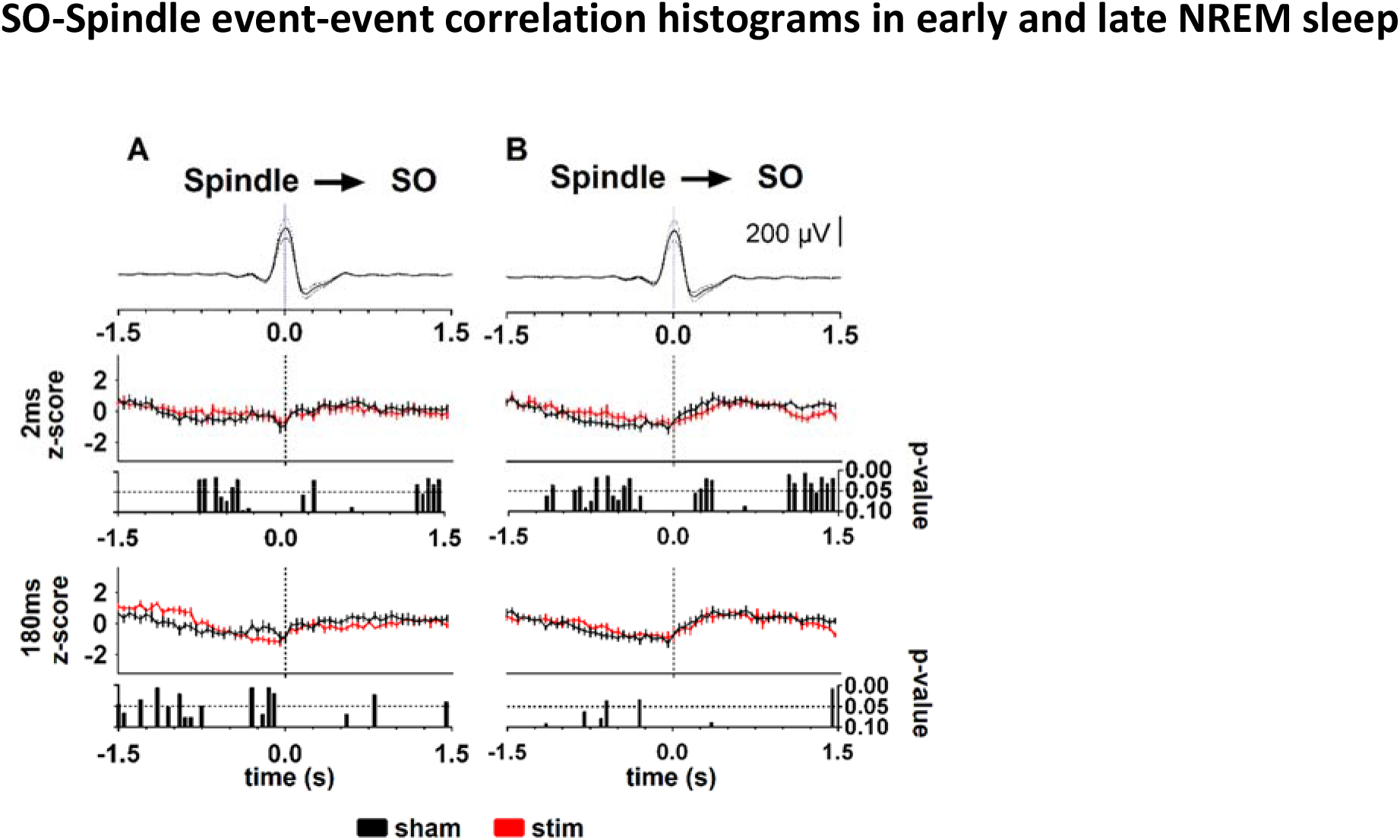
Event-event coupling histograms of spindle activity time-locked to SOs. **A.** Spindle event-activity time-locked to the positive half-wave peak of the SO (t= 0) in early NREM sleep. **B.** Same as A for late NREM sleep. Upper diagrams represent the mean SO event in the sham condition, the bottom bar charts depict the bin-wise p values of two-tailed t-tests (uncorrected for multiple comparisons). n=10, Mean ± SEM.

## Discussion

We aimed to investigate the hippocampal-thalamo-cortical interactions and the corresponding effect on a spatial memory task induced by acoustic stimulation delivered at different SO phases. We found firstly, that of the acoustic stimulation conditions Up-state sACLS, as expected, but also the Down-to-Up-state sACLS revealed a preference index on the OPR task above chance, indicating successful memory for object location. sACLS at the Down state and at the Up-to-Down transition failed to achieve an above chance preference index. The preference index of Up-state sACLS differed significantly from sACLS at a suboptimal SO phase. The lack of significance over sham was most likely due to our short retention interval. Our 3 h retention interval was longer than previously used for successful retention by a group highly experienced with recognition tasks (Dere et al. 2005). A more recent study, while also observing above chance level performance at 2 h, reported a failure of mice to perform the OPR at above chance level at 4 h. A measurement at 3 h was not conducted (Murai et al. 2007).

Secondly, event related activity and stimulus-event correlation histograms both indicate an acute enhancement and subsequent suppression of mean hippocampal SPWR-activity, at all delays. The unique pattern of event related hippocampal activity (cp. also Eschenko et al. 2008) was preserved only for Up-state ACLS. At this phase, the strong enhancement of hippocampal event-activity coincided temporally with the endogenous SO-related increase in hippocampal activity. For sACLS at all other delays, the stimulus-induced increment in hippocampal activity either completely disturbed the endogenous pattern (Down-state sACLS), or evoked activity at an earlier or later time. Our results clearly show that the temporal organization of hippocampal activity and behavior are affected by sACLS phase underscoring a strong connection between intact hippocampal activity during NREM sleep and memory retention on this spatial task. Recent experimental and modelling research reveal increasingly complex neurophysiological interdependencies between cortical and hippocampal activity during NREM sleep (Wei et al. 2020; Azimi et al. 2021; Sanda et al. 2021; Skelin et al. 2021).

We found thirdly that sACLS at all SO phases led to suppressed thalamo-cortical spindle RMS and event-activity. Although human studies with acoustic stimulation support a spindle refractory period (Ngo et al. 2013b; Weigenand et al. 2016; Antony et al. 2018) suppression of spindle activity was not as pronounced, relative to the foregoing enhancement, as observed here. Studies in rodents with a more closely comparable stimulus design to that in humans are therefore warranted. In general, it is however to be considered that deviances in responses may stem from the measurement of local vs. far field potentials, and/or from varying topographical regions, as discussed further below (Khani et al. 2019).

Surprisingly, we did not observe strong modifications by sACLS in event-event coupling despite behavioral differences. At most during early NREM sleep a slight enhancement in modulation of spindle activity by SO activity at Up-state ACLS becomes visible, and modulation of spindle event-activity by SO appeared to be disrupted for all other sACLS conditions. Thus, this slight modulation together with the endogenously coherent enhancement in hippocampal SPWR event-activity the slight modulation in SO-spindle coupling may explain the observed differences in retention performance.

It is generally hypothesized that an increase in coupling strength and/or increased focusing of phase angles coincide with facilitated brain inter-regional communication. Our results point in this direction. However, as recently pointed out by McKillop and colleague, it is important to take not only the global and local properties of sleep into account, which reveal species specific differences, but also recording specificities in interpreting data (McKillop and Vyazovskiy 2020). In the present study cortical activity was recorded from the cingulate cortex. We explicitly did not record from mPFC as in a foregoing study which revealed more pronounced modulation in coupling of SPWRs to SOs (Maingret et al. 2016; Binder et al. 2019; Todorova and Zugaro 2020), since we observed that the shape of (global) SOs varies more with slightly different positioning in the mPFC than cingulate cortex (CG, Binder et al. 2019). Moreover, spindle events were more readily detectable in CG than in the infralimbic mPFC. It could therefore be that due to our choice of CG as cortical recording site hippocampo-cortical connectivity was not optimally reflected. Recording from the ventral hippocampus may also have produced stronger sACLS effects. Although the dorsal hippocampus was recently described to be involved in direct prefrontal to hippocampal communication, monosynaptic hippocampal to prefrontal communication arises from the ventral hippocampus (Binder et al. 2019; Chao et al. 2020). Notably, since mice were successful on the OPR task also in sham, the finding that Up-state sACLS did not produce consistent differences in electrophysiological activity compared to sham is consistent with behavior.

A fourth, unexpected finding was the difference in PreREM sleep duration which was significantly longer at SO-Up state stimulation than for the behaviorally suboptimal sACLS at the SO Up-to-Down state transition. Time spent in all other sleep stages was the same at all sACLS delays. It has been recently reported that spindle count particularly during NREM/ REM sleep transitions which would be scored as PreREM sleep in this study, is a good predictor of behavioral performance in mice during post learning sleep (Yuan et al. 2021). Although, we did not specifically count the density of spindle-like activity during this transition period, our results yield consistently, that PreREM sleep duration was longer at Up-state sACLS than at the later stimulation at 300ms delay. Although results did not uphold multiple comparisons testing NREM sleep spindle duration was also longer at the sACLS Up-state than the Up-to-Down state transition.

In summary, we showed that Down-state sACLS can impair, and Up-state ACLS may potentially enhance performance on the OPR task in mice. Of the measures assessed, SO-related temporal pattern of ripple RMS, stimulus-event-correlations, and event-event-correlations during the first 1-1.5 hours of NREM sleep may most closely explain the behavioral differences of sACLS on memory retention. Our findings support the further use of sACLS for investigating brain responses, changes in inter-regional temporal dynamics, and functional relevance for memory retention.

## Materials and Methods

### Animals

The study was conducted on 10 male C5BL/6N mice (Janvier, France), 8-10 weeks on arrival. Surgery was performed between 10-12 weeks of age. Housing was in Type II-Long IVC cages (Greenline, Tecniplast, GM500) on a 12 h light/dark cycle (lights on: 7:00 h or 8:00 h) with ad libitum access to food and water; initially with littermates and individually after electrode implantation. Animals were handled for 5 min per day for 5 days before the surgery. All animals were treated identically, and all procedures were concordant with the European and national guidelines (EU Directive 2010/63/EU) and were approved by the local state authority (Ministerium für Energiewende, Landwirtschaft, Umwelt und ländliche Räume, Schleswig-Holstein).

### Electrode implantation

Isoflurane (induction: 3.5%, maintenance 0.8-2% at 1-1.3 l/min O2) was used to anaesthetize the animals. They were placed into a stereotactic frame (David Kopf Instruments). To ease the breathing 0.04 mg/kg atropine (Atropinum Sulfuricum, Eifelfango) was administered s.c. After shaving and disinfection of the area, lidocaine (1% solution, B. Braun Melsungen) was injected s.c. before any scalp incision. Every 30 min of surgery 0.1 ml of warm saline was s.c. to substitute fluid loss. An array of five tungsten wires (40 μm, California Fine Wire) was implanted into cingulate gyrus, CG (AP: 1.2, L: 0.3, DV: −0.75) and another array of five tungsten wires, cut slightly diagonally, was implanted into the dorsal hippocampus, HC (AP: −1.94, L: 1.5, DV; deepest wire: −1.35) for recording LFP activity. Two stainless-steel screws (Bilaney, Germany) were used as reference and ground electrodes, implanted above the cerebellum (AP: −4.80, L: 0.00) and the somatosensory cortex (AP: −2, L: 2), respectively. A polyimide-insulated stainless-steel wire (0.125mm diameter, Plastics One) inserted into the neck muscles was used for recording EMG. Wire ends of all electrodes were soldered to a plug-connector during surgery and secured with dental acrylic (Super-Bond, Sun Medical or Relyx Unicem 2 Automix, 3M; Grip Cement, 3M; and Palapress, Heraus Kulzer). Finally, the headstage was encircled by copper wire mesh. Before discontinuing the isoflurane supply carprofen (5 mg/kg, Rimadyl, Pfizer) given i.p. for pain relief. To substitute for fluid loss at the end of the surgery 0.5 ml of warm saline were given s.c. Animals were kept under red light in their home cage until they became mobile and were then transferred back to the animal holding room where they spent at least 7 days for recovery.

### Electrophysiological data acquisition

Recording sessions took place in dark gray recording boxes PVC (20 × 20 × 31 cm; Fachhochschule Lübeck PROJEKT, Germany) with ad libitum access to food and water. The electrode connector was attached to a headstage microamplifier (μPA16, Multi Channel Systems MCS GmbH, Germany). Electrophysiological signals were amplified (amplifier gain: 400) and digitally sampled with 4 kHz through a portable ME16 System (ME16-FAI-μPA, Multi Channel Systems MCS GmbH, Germany) and stored on an assigned computer (MC_Rack software, Multi Channel Systems MCS GmbH, Germany). Real time analog signals from one cortical and one hippocampal electrode were transferred through two audio outputs of the ME16 System to the data acquisition interface CED Power1401-3 (Cambridge Electronic Design Limited, Cambridge, England) which was connected to a second computer. The CED 1401-3, its application program Spike 2 and a custom-made program based on the built-in script language were used for online calculation of the hippocampal delta/theta ratio and the cortical SO detection.

### Threshold determination for the single acoustic closed loop stimulation during NREM sleep

The second habituation recording was used for offline determination of the hippocampal delta/theta ratio threshold, which was used in the experimental recording sessions for online NREM sleep detection and to determine the initial threshold for the cortical signal that was to be used for online SO detection for each individual animal. The threshold of the delta/theta ratio was set by visual inspection from two histograms showing the distribution of all delta/theta ratio values of all sleep stages, one histogram for NREM sleep and another histogram for all other sleep stages (Wake, REM, PreREM, see Supplementary Figure 4). Analyses to determine the SO detection threshold were limited to NREM sleep epochs, and the same SO detection algorithm used for all offline SO analyses, was applied. This algorithm is described in detail further below.

### Single acoustic closed loop stimulation

An acoustic stimulation was given or omitted (‘pseudo-stimulation’) upon detection of a positive potential of the cortical signal superseding the SO detection threshold when the delta/theta ratio was above the threshold for NREM sleep unless the detection algorithm was paused for some reason by the experimenter. Following SO detection and corresponding stimulation, the algorithm was paused automatically for 2.5 s. A trigger devoid of stimulation was generated for every other detected SO; i.e. acoustic stimulation and pseudo-stimulation alternated. For acoustic stimulation an analog signal, consisting of 10 ms acoustic white noise, was sent from the Power1401-3 to two dome tweeters (TW 6 NG 8 ohm, Visaton, Germany) inside of the recording box through a custom-made stereo audio amplifier (2 x 5 W) at 58 dBA volume. Online detection of SOs was conducted by a custom-made script running under Spike 2 software together with a sequencer in the Power1401-3, like that described in Ngo et al (2013). In brief, each time the EEG signal crossed an adaptive threshold toward larger positive values, the acoustic stimulation was triggered. On default, the threshold was set to the initial value calculated from the second habituation recording. Every 0.5 s, the threshold was updated to the maximal (i.e., largest positive) instantaneous amplitude of the cortical signal within the preceding 2 s interval, however, only if this value exceeded the initial threshold. This algorithm ensured a continuously reliable detection of gradually increasing and decreasing SO amplitude during sleep or within a sleep cycle by its positive half-wave peak.

The timing of all SO detections was marked digitally (with different numbers for acoustic stimulation and pseudo-stimulation) in the Spike 2 data file and these markers were also transferred to the ME16 System to be saved with the MC_Rack Software along with the animals cortical and hippocampal data on a separate channel. The delay between the SO detection and the delivery of the stimulation differed between each stimulation condition, e.g.; 2 ms, 120 ms, 180 ms, 300 ms. The fifth condition was the sham condition in which the acoustic output was omitted during the entire recording. Each condition was applied on a separate day and separated by about one week.

## Behavioral procedures

### Habituation to the recording procedure

After the post-surgery recovery period, animals were habituated to the behavioral open field and recording boxes. Prior to each recording habituation animals were transferred to the experimental room before the end of the last light phase, to spend the subsequent dark phase in the recording box, untethered. After lights on, mice were connected to the head stage and electrophysiological activity was monitored for 3h. Habituation recordings took place on two consecutive days and the data from the second day was analyzed to determine parameters for online SO detection and delta/theta ratio.

### Habituation to the object place recognition task

Animals were brought to the experimental room 1h prior to the habituation and at first left to adapt to the room in their home cages. Light levels were set to 100 lux above the open field for all experiments. After adaptation, mice were placed in to the middle of the open field (37×37×35 cm, Ewald Kongsbak GmbH + Co. KG) via a transfer cage and allowed to explore the field for 10 minutes. This procedure took place daily on 3 consecutive days prior to the OPR task.

### Object place recognition task

All OPR tasks took place at the beginning of the light phase. The mean age of the animals on the first day of the OPR was 97.90 ± 6.15 days and 137.30 ± 9.55 days on the last day. Animals were transferred to the open field as during the habituation. In brief, in the OPR sample phase (10 min) mice were exposed to two identical objects, which were placed in two corners of the field. Special care was taken that the surface of the objects that animals contacted would not differ. For the test phase (5 min) one of the objects was then displaced to one of the other two free corners whereas the other object remained at its position. Both objects were cleaned with 70% ethanol before each sample and test phase, as well as after each OPR session. Electrophysiological recording took place within the 3 h-interval between the two OPR phases.

Each animal participated in 5 OPR trials corresponding to the 5 conditions (4 different delay times after the SO Down state for stimulation, and sham). The OPR sample phase started on average 39.2 ± 2.0 min and electrophysiological recording 55.4 ± 2.1 min after lights on. OPR sessions were set 5-8 days apart in order to prevent interference from previous memories. Positioning of the displaced object was pseudo-randomized.

## Data analysis

### Object place recognition task

Video-recorded behavior was scored manually by an experimenter blind to the experimental condition and previous placement of the objects using a tracking software (AnyMaze, Stoelting, version 4.72). The time spent by mice actively exploring each object was rated manually. Grooming or freezing behavior in close vicinity to objects were not included. Spatial retention performance was measured as the preference index, calculated as the ratio of exploration time of the displaced object to the total exploration time of both objects during the test phase.

### Sleep Architecture

An experimental rater assigned sleep stages to all 5 s epochs of the 3h sleep recording according to the standard criteria using the cortical and hippocampal LFP as well as EMG recordings (SleepSign for Animals, Kissei Comtec). Briefly, the sleep stages were characterized as follows: wakefulness (Wake) by high EMG activity and desynchronized CG LFP; NREM sleep by low EMG and high-amplitude low frequency CG LFP activity consisting mostly of delta activity (0.75-4 Hz); pre-rapid eye movement sleep (PreREM) by low EMG and high-amplitude CG sigma (10-15 Hz) activity before REM sleep or Wake; and REM sleep by low EMG activity and high theta activity (6-10 Hz) in the HC LFP.

Furthermore, after scoring the 3 h sleep period, NREM sleep was divided into two equal length periods, without splitting sleep cycles, termed ‘early’ and ‘late’ NREM sleep, to compare selected effects.

### Electrophysiology

Electrophysiological analyses were conducted on the LFP recording from the NREM sleep epochs over the 3 h post-learning period or within early and late NREM sleep periods. LFP data included one chosen CG and HC channel from each animal. Data were analyzed using Spike2 (Cambridge Electronic Design) and custom scripts written with the built-in script language.

### Offline event detection

SO, spindle and ripple events were identified similar to Binder et al. (2019). In brief, to identify SOs in the CG LFP signal, a low-pass finite impulse response (FIR) filter of 30 Hz was applied and the resultant signal was down-sampled to 100 Hz. Subsequently, a low-pass FIR filter of 3.5 Hz was used to produce the SO signal. In the slow oscillation signal all two succeeding negative-to-positive zero crossings separated by 0.45–1.43 s (corresponding to 0.7–2.22 Hz) were marked and the negative and the positive peak potentials between these marked negative-to-positive zero crossings were registered. SO events were defined as those intervals that displayed (1) a positive peak amplitude of 1.25 times the average positive peak amplitude of this animals sham session or higher (2) a positive-to-negative peak amplitude difference of at least 1.25 times the average positive-to-negative peak amplitude difference of the respective sham session.

Spindle identification also required first low-pass (<30 Hz) filtering and down-sampling to 100 Hz of the CG LFP. Subsequently a FIR bandpass filter of 9 – 15 Hz was applied and a root mean square (RMS) representation of the filtered signal was generated by using a 0.2 s long sliding window. The resulting RMS signal was smoothed additionally with a sliding window average of 0.2 s. Time frames were considered as spindle intervals if the RMS signal exceeded a threshold of 1.25 SD of the bandpass filtered signal for 0.5 – 3 s and if the largest value within the frame was >2 SD of the bandpass filtered signal. For each animal individual thresholds were calculated from the bandpass filtered signal of the sham session. Two succeeding spindles were counted as one spindle when the interval between the end of the first spindle and the beginning of the second spindle was shorter than 0.5 s and the resulting (merged) spindle was not >3 s. Detected events were not accepted as spindles, when the difference between the largest and smallest potential of the low-pass filtered signal (<30 Hz) within the frame was 5 times larger than 2 SD of the bandpass filtered signal and the time between these two extrema was equal or shorter than one-half of an oscillation cycle of 15 Hz (0.033 s). For event-correlation analyses the peaks and troughs of every spindle were marked as the maxima and minima of the bandpass filtered signal (between the beginning and end of the spindle), and the deepest trough was designated as the “spindle peak” that represented the respective spindle in time, i.e., the time point taken for referencing event correlation histograms (see description of event correlation histograms later in this section).

Identification of ripples initially required the application of a low-pass FIR filter of 300 Hz and down-sampling to 1 kHz of the dorsal hippocampal LFP. Subsequently, a bandpass FIR filter of 150 – 200 Hz was applied and the RMS signal was calculated with a sliding 0.02 s time window. The RMS signal was then smoothed with a sliding window average of 0.02 s. Time frames were considered as ripple intervals if the RMS signal exceeded a threshold of 1.25 SD of the bandpass filtered signal for 0.025 – 0.1 s and if the largest value within the frame was >5 SD of the bandpass filtered signal. For each animal individual thresholds were calculated from the bandpass filtered signal of the sham session. Detected events were not accepted as ripples, when the difference between the largest and smallest potential of the low-pass filtered signal (<300 Hz) within the frame was 5 times larger than 5 SD of the bandpass filtered signal and the time between these two extrema was equal or shorter than one-half of an oscillation cycle of 200 Hz (2.5 ms). For event-correlation analyses, the peaks and troughs of every ripple were marked as the maxima and minima of the bandpass filtered signal, and the deepest trough was designated as the “ripple peak” that represented the respective ripple in time.

Durations of SO, spindle and SPWR oscillatory events correspond to the time from beginning to end of the detected oscillatory event, respectively. Density values refer to the mean density of the corresponding event across all NREM sleep epochs. SO, spindle and ripple amplitudes are defined as the mean peak-to-peak values of events detected from the corresponding filtered frequency band signal. Spindle and ripple RMS represent the average values across all detected oscillatory events. The SO slope was derived from the positive Down state peak to the following zero crossing of the SO filtered signal. Grand mean averages of the detected SOs, spindles, and ripples across all animals were calculated for the different conditions. Also, averages of the original signal, spindle RMS and ripple RMS were calculated for acoustic stimulation of all delays and corresponding pseudo-stimulation.

### Stimulus-event correlation histograms

Stimulus-event correlation histograms were calculated for SO (positive half-wave peaks), spindle and ripple activity (number of peaks and troughs) in intervals of −1.0 — 2.5 s around the online detected and stimulated SOs. These histograms, with a bin size of 50 ms, were separately calculated for all acoustic- and all pseudo-stimulations during NREM sleep over the entire 3 h recording session. The individual histograms were z-scored by the corresponding mean and SD of the SO, spindle and ripple activity, respectively, for each animal during the −1.0 — 2.5 s interval to eliminate the considerable variability across animals and conditions. These histograms represent the acute modulatory effect of sACLS on event activity. Grand mean averages of the stimulus-event correlation histograms across all animals were calculated for the different conditions.

### Event-event correlation histograms

Event-event correlation histograms were calculated for spindle and ripple activity (number of peaks and troughs) with reference to the time of the SO positive half-wave peak as identified in the CG LFP signal. Further event-event correlation histograms were calculated for ripple activity (number of peaks and troughs) in the dorsal hippocampal LFP with reference to the spindles (i.e., spindle peaks, see description of spindle identification in the prior paragraph) and for spindle activity (number of peaks and troughs) in the CG LFP with reference to the ripples (i.e., ripple peak, see description of ripple identification in the prior paragraph). For all event-event correlation histograms, 3 s windows were used with an offset of 1.5 s and a bin size of 50 ms. Again, these histograms were calculated for all NREM sleep epochs over the entire 3 h recording session and additionally for early and late NREM sleep. The individual histograms were z-scored by the corresponding mean and SD of the spindle and ripple activity, respectively, for each animal during the ±1.5 s interval to eliminate the considerable variability across animals and conditions. In contrast to the stimulus-event correlation histograms, the event-event correlation histograms were independent of the timing of sACLS and compared the sham session to the other stimulation sessions. The histograms represent a measure for the probability of activity of one event at a given time to proceed or follow another event, i.e., for the coupling of SOs, cortical spindles and hippocampal ripples.

### Experimental Design and Statistical Analysis

The study was conducted in a within-subject design, with 5 sessions. For statistical analyses of the preference index the mean score of each session was first compared to the odds ratio of 0.5 by a one sample t-test. Behavioral data were then subjected to a mixed effects model for repeated measures analyses of variance and post-hoc paired t-tests.

Event parameters of SOs, spindles, and SPWRs; duration of sleep stages; and number of stimulation triggers given at a certain sleep stage for the 3 h interval or for early and late NREM sleep periods were subjected to mixed effects models for repeated measures with the factors Condition (sham, 2 ms, 120 ms, 180 ms, 300 ms delay), and when applicable Time (Early, Late). Stimulus-event histograms of acoustic and pseudo-stimulation were compared by paired t-tests with correction for multiple comparisons using the Holm-Sidak method. Comparisons between delay conditions and Sham in the event-event correlation histograms (over the 3h period and within early and late periods) were similarly conducted by paired t-tests, but not corrected for multiple comparisons. All data are expressed as mean ± SEM and analyzed using the software package Prism 8 (GraphPad Software, Inc.). A p value < 0.05 was considered significant.

## Acknowledgements

We gratefully thank Erik Gromodka for preparing the figures, and Sonja Binder for technical support. This work was supported by the US-German Collaboration in Computational Neuroscience (NSF/BMBF grant 01GQ1706) and IIS-1724405), National Institute of Health (grants 1RO1MH125557 and 1RO1MH117155).

## Notes

### Competing Interest Statement

The authors have declared no competing interest.

## References

Antony JW, Piloto L, Wang M, Pacheco P, Norman KA, Paller KA. 2018. Sleep Spindle Refractoriness Segregates Periods of Memory Reactivation. Curr Biol 28: 1736–1743.e1734.

Azimi A, Alizadeh Z, Ghorbani M. 2021. The essential role of hippocampo-cortical connections in temporal coordination of spindles and ripples. Neuroimage 243: 118485.

Barker GR, Warburton EC. 2011. When is the hippocampus involved in recognition memory? J Neurosci 31: 10721–10731.

Binder S, Mölle M, Lippert M, Bruder R, Aksamaz S, Ohl F, Wiegert JS, Marshall L. 2019. Monosynaptic Hippocampal-Prefrontal Projections Contribute to Spatial Memory Consolidation in Mice. J Neurosci 39: 6978–6991.

Campos-Beltran D, Marshall L. 2017. Electric Stimulation to Improve Memory Consolidation During Sleep. In Cognitive Neuroscience of Memory Consolidation (ed. N Axmacher, B Rasch), pp. 301–312. Springer.

Chao OY, de Souza Silva MA, Yang YM, Huston JP. 2020. The medial prefrontal cortex - hippocampus circuit that integrates information of object, place and time to construct episodic memory in rodents: Behavioral, anatomical and neurochemical properties. Neurosci Biobehav Rev 113: 373–407.

Cromwell HC, Mears RP, Wan L, Boutros NN. 2008. Sensory gating: a translational effort from basic to clinical science. Clin EEG Neurosci 39: 69–72.

Dere E, Huston JP, De Souza Silva MA. 2005. Episodic-like memory in mice: simultaneous assessment of object, place and temporal order memory. Brain Res Brain Res Protoc 16: 10–19.

Ego-Stengel V, Wilson MA. 2010. Disruption of ripple-associated hippocampal activity during rest impairs spatial learning in the rat. Hippocampus 20: 1–10.

Eschenko O, Mölle M, Born J, Sara SJ. 2006. Elevated sleep spindle density after learning or after retrieval in rats. J Neurosci 26: 12914–12920.

Eschenko O, Ramadan W, Mölle M, Born J, Sara SJ. 2008. Sustained increase in hippocampal sharp-wave ripple activity during slow-wave sleep after learning. Learn Mem 15: 222–228.

Gais S, Mölle M, Helms K, Born J. 2002. Learning-dependent increases in sleep spindle density. J Neurosci 22: 6830–6834.

Girardeau G, Benchenane K, Wiener SI, Buzsáki G, Zugaro MB. 2009. Selective suppression of hippocampal ripples impairs spatial memory. Nature Neuroscience 12: 1222–1223.

Giri B, Miyawaki H, Mizuseki K, Cheng S, Diba K. 2019. Hippocampal Reactivation Extends for Several Hours Following Novel Experience. The Journal of Neuroscience 39: 866–875.

Harrington MO, Ngo HV, Cairney SA. 2021. No benefit of auditory closed-loop stimulation on memory for semantically-incongruent associations. Neurobiol Learn Mem 183: 107482.

Helfrich RF, Lendner JD, Mander BA, Guillen H, Paff M, Mnatsakanyan L, Vadera S, Walker MP, Lin JJ, Knight RT. 2019. Bidirectional prefrontal-hippocampal dynamics organize information transfer during sleep in humans. Nature Communications 10.

Henin S, Borges H, Shankar A, Sarac C, Melloni L, Friedman D, Flinker A, Parra LC, Buzsaki G, Devinsky O et al. 2019. Closed-Loop Acoustic Stimulation Enhances Sleep Oscillations But Not Memory Performance. eneuro 6: ENEURO.0306-0319.

Ketz N, Jones AP, Bryant NB, Clark VP, Pilly PK. 2018. Closed-Loop Slow-Wave tACS Improves Sleep-Dependent Long-Term Memory Generalization by Modulating Endogenous Oscillations. J Neurosci 38: 7314–7326.

Khani A, Lanz F, Loquet G, Schaller K, Michel C, Quairiaux C. 2019. Large-Scale Networks for Auditory Sensory Gating in the Awake Mouse. eneuro 6.

Ladenbauer J, Ladenbauer J, Külzow N, Flöel A. 2021. Memory-relevant nap sleep physiology in healthy and pathological aging. Sleep 44.

Maingret N, Girardeau G, Todorova R, Goutierre M, Zugaro M. 2016. Hippocampo-cortical coupling mediates memory consolidation during sleep. Nature Neuroscience 19: 959–964.

Malkani RG, Zee PC. 2020. Brain Stimulation for Improving Sleep and Memory. Sleep Med Clin 15: 101–115.

Marshall L, Helgadóttir H, Mölle M, Born J. 2006. Boosting slow oscillations during sleep potentiates memory. Nature 444: 610–613.

McKillop LE, Vyazovskiy VV. 2020. Sleep and ageing: from human studies to rodent models. Curr Opin Physiol 15: 210–216.

Moreira CG, Baumann CR, Scandella M, Nemirovsky SI, Leach S, Huber R, Noain D. 2021. Closed-loop auditory stimulation method to modulate sleep slow waves and motor learning performance in rats. eLife 10.

Mumby DG, Gaskin S, Glenn MJ, Schramek TE, Lehmann H. 2002. Hippocampal Damage and Exploratory Preferences in Rats: Memory for Objects, Places, and Contexts. Learning & Memory 9: 49–57.

Murai T, Okuda S, Tanaka T, Ohta H. 2007. Characteristics of object location memory in mice: Behavioral and pharmacological studies. Physiol Behav 90: 116–124.

Ngo HV, Claussen JC, Born J, Mölle M. 2013a. Induction of slow oscillations by rhythmic acoustic stimulation. J Sleep Res 22: 22–31.

Ngo HV, Martinetz T, Born J, Mölle M. 2013b. Auditory closed-loop stimulation of the sleep slow oscillation enhances memory. Neuron 78: 545–553.

Ngo HV, Miedema A, Faude I, Martinetz T, Mölle M, Born J. 2015. Driving sleep slow oscillations by auditory closed-loop stimulation-a self-limiting process. J Neurosci 35: 6630–6638.

Novitskaya Y, Sara SJ, Logothetis NK, Eschenko O. 2016. Ripple-triggered stimulation of the locus coeruleus during post-learning sleep disrupts ripple/spindle coupling and impairs memory consolidation. Learning & Memory 23: 238–248.

Ong JL, Lo JC, Chee NI, Santostasi G, Paller KA, Zee PC, Chee MW. 2016. Effects of phase-locked acoustic stimulation during a nap on EEG spectra and declarative memory consolidation. Sleep Med 20: 88–97.

Oyanedel CN, Durán E, Niethard N, Inostroza M, Born J. 2020. Temporal associations between sleep slow oscillations, spindles and ripples. European Journal of Neuroscience 52: 4762–4778.

Rasch B, Born J. 2013. About sleep’s role in memory. Physiol Rev 93: 681–766.

Salfi F, D’Atri A, Tempesta D, De Gennaro L, Ferrara M. 2020. Boosting Slow Oscillations during Sleep to Improve Memory Function in Elderly People: A Review of the Literature. Brain Sci 10.

Sanda P, Malerba P, Jiang X, Krishnan GP, Gonzalez-Martinez J, Halgren E, Bazhenov M. 2021. Bidirectional Interaction of Hippocampal Ripples and Cortical Slow Waves Leads to Coordinated Spiking Activity During NREM Sleep. Cereb Cortex 31: 324–340.

Sawangjit A, Oyanedel CN, Niethard N, Salazar C, Born J, Inostroza M. 2018. The hippocampus is crucial for forming non-hippocampal long-term memory during sleep. Nature 564: 109–113.

Schapiro AC, Reid AG, Morgan A, Manoach DS, Verfaellie M, Stickgold R. 2019. The hippocampus is necessary for the consolidation of a task that does not require the hippocampus for initial learning. Hippocampus 29: 1091–1100.

Schneider J, Lewis PA, Koester D, Born J, Ngo H-VV. 2020. Susceptibility to auditory closed-loop stimulation of sleep slow oscillations changes with age. Sleep 43.

Skelin I, Zhang H, Zheng J, Ma S, Mander BA, Kim McManus O, Vadera S, Knight RT, McNaughton BL, Lin JJ. 2021. Coupling between slow waves and sharp-wave ripples engages distributed neural activity during sleep in humans. Proc Natl Acad Sci U S A 118.

Stickgold R. 2005. Sleep-dependent memory consolidation. Nature 437: 1272–1278.

Süer C, Dolu N, Ozesmi C. 2004. The effect of immobilization stress on sensory gating in mice. Int J Neurosci 114: 55–65.

Swift KM, Gross BA, Frazer MA, Bauer DS, Clark KJD, Vazey EM, Aston-Jones G, Li Y, Pickering AE, Sara SJ et al. 2018. Abnormal Locus Coeruleus Sleep Activity Alters Sleep Signatures of Memory Consolidation and Impairs Place Cell Stability and Spatial Memory. Current Biology 28: 3599–3609.e3594.

Todorova R, Zugaro M. 2020. Hippocampal ripples as a mode of communication with cortical and subcortical areas. Hippocampus 30: 39–49.

Tononi G, Cirelli C. 2014. Sleep and the Price of Plasticity: From Synaptic and Cellular Homeostasis to Memory Consolidation and Integration. Neuron 81: 12–34.

Wei Y, Krishnan GP, Marshall L, Martinetz T, Bazhenov M. 2020. Stimulation Augments Spike Sequence Replay and Memory Consolidation during Slow-Wave Sleep. J Neurosci 40: 811–824.

Weigenand A, Mölle M, Werner F, Martinetz T, Marshall L. 2016. Timing matters: open-loop stimulation does not improve overnight consolidation of word pairs in humans. Eur J Neurosci 44: 2357–2368.

Witten L, Bastlund JF, Glenthøj BY, Bundgaard C, Steiniger-Brach B, Mørk A, Oranje B. 2016. Comparing Pharmacological Modulation of Sensory Gating in Healthy Humans and Rats: The Effects of Reboxetine and Haloperidol. Neuropsychopharmacology 41: 638–645.

Yuan RK, Lopez MR, Ramos-Alvarez M-M, Normandin ME, Thomas AS, Uygun DS, Cerda VR, Grenier AE, Wood MT, Gagliardi CM et al. 2021. Differential effect of sleep deprivation on place cell representations, sleep architecture, and memory in young and old mice. Cell Reports 35: 109234.

