## Supplemental Figures for "Single acoustic closed loop stimulation in mice to modulate hippocampo-thalamo-cortical activity and performance"

Supplemental_Fig_S1.pdf

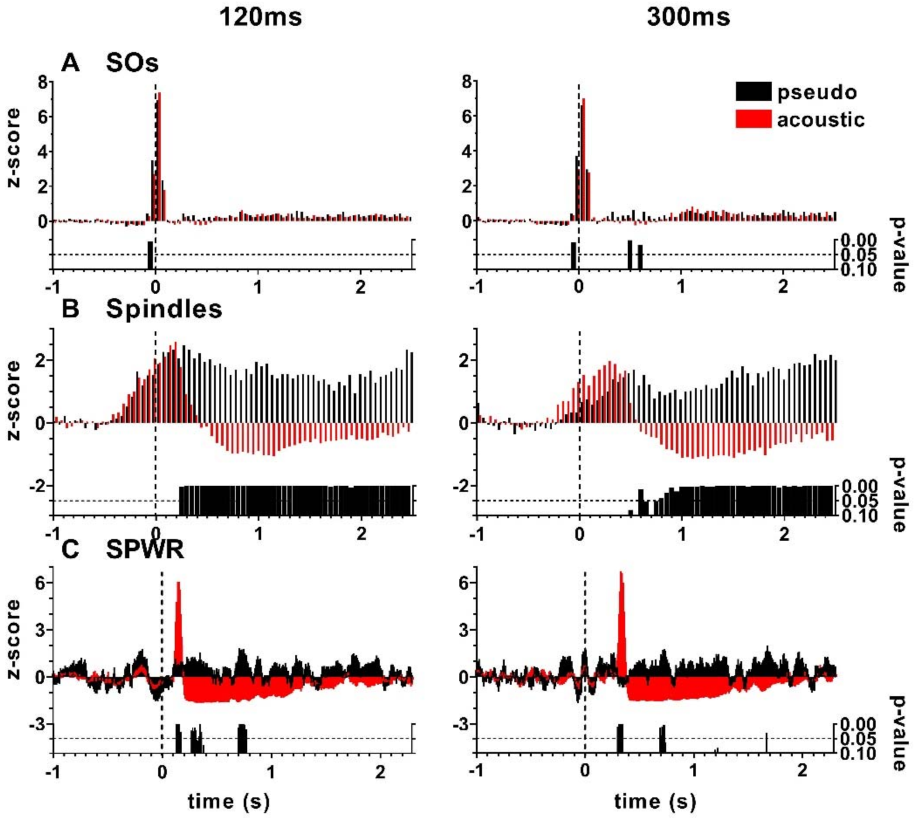


**Supplemental Figure 1. SO , thalamo-cortical spindle and hippocampal ripple event-activity relative to sACLS. A-C.** Mean stimulus-event correlations for SO, spindle and CA1 SPWR events (z-transformed) for acoustic and pseudo-stimulation averaged across the 3h interval for the 120 ms and 300 ms delay conditions, respectively. T = 0 corresponds to the detected SO Down state peak. Data are baseline normalized across the first 0.5 seconds. Bar charts depict the Sidak-Holms adjusted bin-wise p-values of two-tailed t-tests. 120ms, n = 7; 300ms, n = 6.

Supplemental_Fig_S2.pdf

**Event-event correlation histograms**

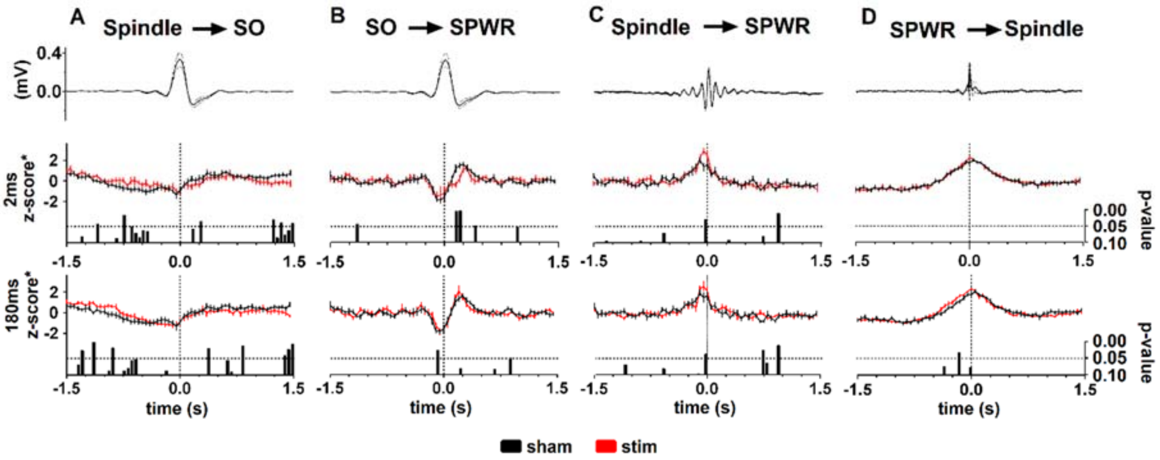


**Supplemental Figure 2. Event-coupling histograms between SO, spindle and SPWR event activity, respectively, across the 3h interval. A.** Spindle event-activity time-locked to the negative half-wave peak of the SO (t= 0). **B.** SPWR event-activity time-locked to the negative half-wave peak of the SO (t= 0). **C.** Ripple event-activity time-locked to the deepest trough of the cortical spindle. **D.** Spindle event-activity time-locked to the deepest trough of hippocampal ripples. **A-D.** Upper diagrams represent the mean reference event in sham condition, bottom bar charts depict the bin-wise p values of two-tailed t-tests (not corrected for multiple comparisons). Mean ± SEM. 120ms, n = 7; 300ms, n = 6.

Supplemental_Fig_S3.pdf

**SO-Spindle event-event correlation histograms in early and late NREM sleep**

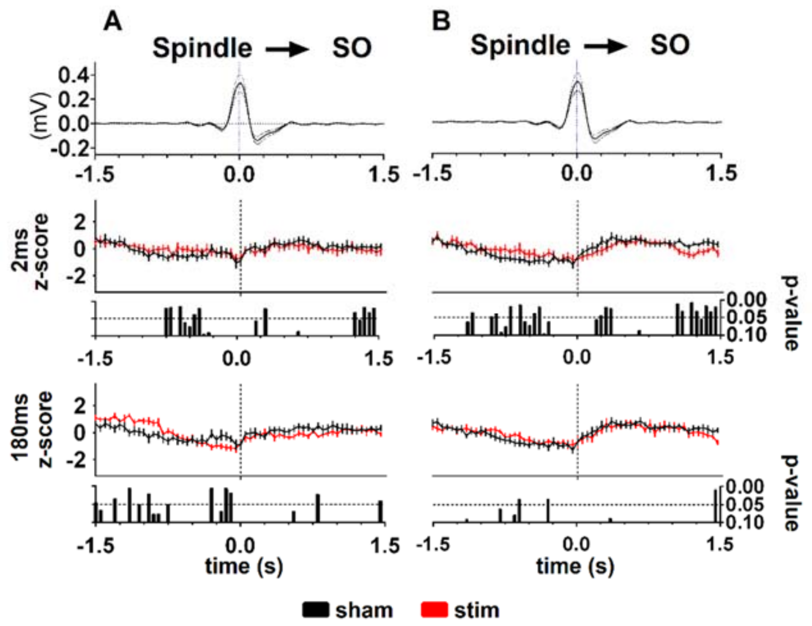


**Supplemental Figure 3 Event-event coupling histograms of spindle activity time-locked to SOs. A.** Spindle event-activity time-locked to the negative half-wave peak of the SO (t= 0) in early NREM sleep. **B.** Same as A for late NREM sleep. Upper diagrams represent the mean SO event in the sham condition, the bottom bar charts depict the bin-wise p values of two-tailed t-tests (uncorrected for multiple comparisons). Mean ± SEM.120ms, n = 7; 300ms, n = 6.

Supplemental_Fig_S4.pdf

**Methods**

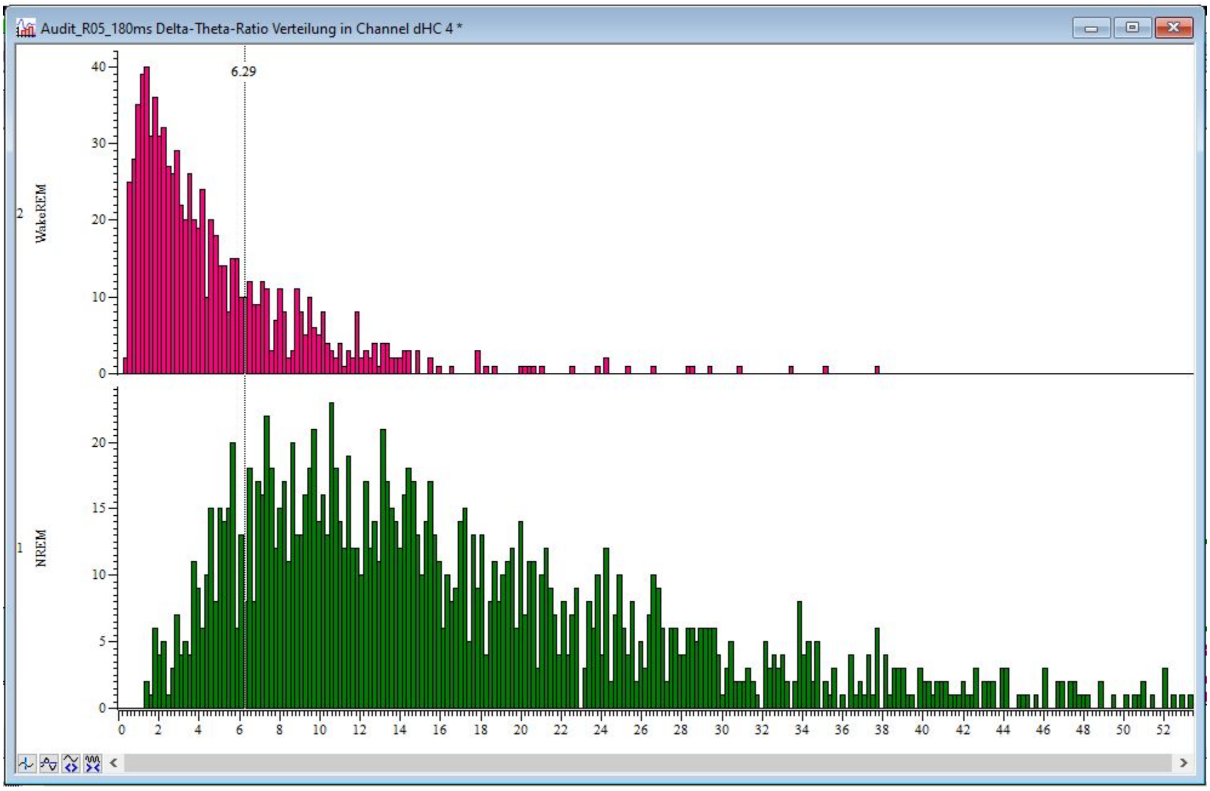


**Supplemental Figure 4. Histograms to determine the delta/theta ratio from a hippocampal LFP recording.** The top histogram gives the number of 10-sec epochs scored as Wake or REM sleep for each delta/theta ratio given on the x-axis. The corresponding bottom histogram is for NREM sleep epochs. The threshold of 6.29 (vertical black bar) indicates the value after which the greater majority of NREM sleep epochs, but minority of wake and REM sleep epochs were scored.
